# BRANE Cut: Biologically-Related A priori Network Enhancement with Graph cuts for Gene Regulatory Network Inference

**DOI:** 10.1101/032383

**Authors:** Aurélie Pirayre, Camille Couprie, Frédérique Bidard, Laurent Duval, Jean-Christophe Pesquet

## Abstract

**Background:** Inferring gene networks from high-throughput data constitutes an important step in the discovery of relevant regulatory relationships in organism cells. Despite the large number of available Gene Regulatory Network inference methods, the problem remains challenging: the underdetermination in the space of possible solutions requires additional constraints that incorporate a priori information on gene interactions.

**Methods:** Weighting all possible pairwise gene relationships by a probability of edge presence, we formulate the regulatory network inference as a discrete variational problem on graphs. We enforce biologically plausible coupling between groups and types of genes by minimizing an edge labeling functional coding for a priori structures. The optimization is carried out with Graph cuts, an approach popular in image processing and computer vision. We compare the inferred regulatory networks to results achieved by the mutual-information-based Context Likelihood of Relatedness (CLR) method and by the state-of-the-art GENIE3, winner of the DREAM4 multifactorial challenge.

**Results:** Our BRANE Cut approach infers more accurately the five DREAM4 in silico networks (with improvements from 6% to 11%). On a real *Escherichia coli* compendium, an improvement of 11.8% compared to CLR and 3% compared to GENIE3 is obtained in terms of Area Under Precision-Recall curve. Up to 48 additional verified interactions are obtained over GENIE3 for a given precision. On this dataset involving 4345 genes, our method achieves a performance similar to that of GENIE3, while being more than seven times faster. The BRANE Cut code is available at: http://www-syscom.univ-mlv.fr/~pirayre/Codes-GRN-BRANE-cut.html

**Conclusions:** BRANE Cut is a weighted graph thresholding method. Using biologically sound penalties and data-driven parameters, it improves three state-of-the-art GRN inference methods. It is applicable as a generic network inference post-processing, due its computational efficiency.

## Background

Gene expression microarray techniques and high-throughput sequencing-based experiments furnish numerical data for gene regulatory process characterization. Gene Regulatory Network (GRN) inference provides a framework to transform high-throughput data into meaningful information. It consists of the construction of graph structures that highlight regulatory links between transcription factors and their target genes. GRNs are used as an initial step for experimental condition analysis or network interpretation, for instance classification tasks [1], leading to more insightful biological knowledge extraction. It may also directly offer genetic targets for specific experiments, such as directed mutagenesis and/or knock-out procedures.

Despite the large variety of proposed GRN inference methods, building a GRN remains a challenging task due to the nature of gene expression and the structure of the experimental data. It notably involves data dimensionality, especially in terms of gene/replicate/condition proportions. Indeed, gene expression data obtained from microarrays or high-throughput technologies correspond to the expression profiles of thousands of genes. Expression profiles reflect gene expression levels for different replicates or strains studied in different physico-chemical, temporal or culture medium conditions. Although the cost of biological experiments diminishes, gene expression data is often acquired under a limited number of replicates and conditions compared to the number of genes. This causes difficulties in properly inferring gene regulatory networks and in recovering reliable biological interpretations of such networks. Continuous efforts from the bioinformatics community, partly driven by the organization of the DREAM challenges [2], hitherto allowed for constant progresses in GRN inference efficiency.

GRN inference approaches are often cleaved into two classes of methods [3, 4]: model-based or information-theoretic score-based. The latter notably employs mutual-information measures, which quantify the mutual dependence or the information shared by stochastic phenomena. They are used in frequently mentioned and compared GRN methods, for instance: Relevance Network (RN) [5], Algorithm for the Reconstruction of Accurate Cellular Network (ARACNE) [6], Minimum Redundancy NETwork (MRNET) [7], or Context Likelihood of Relatedness (CLR) [8]. CLR was shown to outperform RN, ARACNE and MRNET on several datasets [8]. While RN removes edges whose mutual information is lower than a threshold, CLR exhibits improved performance by computing a score derived from *Z*-statistics on mutual-information, leading to more robust results. Model-based methods include Bayesian approaches, Gaussian graphical models [9, 10], or differential equations [11, 4]. Graphical models rely on strong hypotheses on data distribution, that may yield poor performance when tested on real datasets where the number of replicates or conditions is very small in proportion to the number of genes. The performance of such inferred networks can be sensibly improved by a network deconvolution approach ([12], thereafter denoted by ND) that removes global transitive or indirect effects by eigen-decompositions. Differential equation approaches are often restricted by limited-size time course data. The more recent GENIE3 (GEne Network Inference with Ensemble of trees) [13] and Jump3 [14] approaches prevent such a pitfall by avoiding assumptions on the data. Instead, they formulate the graph inference as a feature selection problem, and learn a ranking of edge presence probability. A drawback of model-based versus mutual-information-based approaches is a rather high computational cost on standard-size networks.

The problem of network inference boils down to finding a set of edges that (hopefully) represents actual regulations between genes, given their expression profiles. As we search for a set of regulatory edges, the outcome can be related to an integer binary solution: presence or absence for each gene-to-gene edge. From this framework, we incorporate additional structural a priori based on biological observations and assumptions. They control different connectivity aspects involving particular genes coding for transcription factors. Such supplementary information from heterogeneous data sources, when available, supports the network inference process [15]. We then translate the network inference problem into a variational formulation as detailed in the ‘Mathematical modeling of the structural a priori’ section. Our approach generalizes classical inference. A first additional penalty influences the degree of connectivity of transcription factors and target genes. A second constraint promotes edges related to co-regulation mechanisms. The obtained integer programming problem may be solved by finding a maximal flow in a graph, as explained in the ‘Optimization strategy’ section. This approach, known as Graph cuts, is well-investigated in the computer vision and image processing literature, where it has demonstrated computational efficiency in a large number of tasks [16].

Our contributions are the following:

1. We introduce BRANE Cut, a novel Biologically-Related A priori Network Enhancement for gene regulation based on Graph cuts. Previous Graph cuts formulations in bioinformatics were employed only for clustering in biological network analysis [17] or for feature selection in the Genome-Wide Association Study context [18].
2. The proposed method generalizes standard regulatory network inference by incorporating additional terms with biological interpretation. Since their regularization parameters are estimated from gene set cardinality, it can be applied to various transcriptomic data.
3. The computation time of our method is negligible in comparison with other model-based approaches with inference improvements.
4. It can be used as a generic GRN post-processing with any input weights and supplementary information on transcription factors.

The paper is organized as follows: we propose in the next section the novel variational approach for building GRNs and we detail the efficient optimization strategy used to solve the related minimization problem. BRANE Cut outcomes and performance on benchmark datasets coming from the DREAM4 challenge and the *Escherichia coli* compendium are provided in the ‘Results and discussion’ section. We finally conclude and offer perspectives.

## Methods

### Mathematical modeling of the structural a priori

We first introduce our notations before detailing our structural models and variational formulation. Let *G* represent the total number of genes for which expression data is collected. Expression data is gathered in a symmetric weighted adjacency matrix **W** ∈ ℝ*^G^*^×^*^G^*. Its (*i,j*) element corresponds to a statistical measure reflecting the strength of the link, or information shared, between the expression profiles of gene *i* ∈ {1, …, *G*} and gene *j* ∈ {1, …, *G*}. Our approach uses nonnegative weights. A convenient choice for *ω_i,j_* is the normalized mutual information (*ω_i,j_* ∈ [0,1]) computed between the expression profiles of genes *i* and *j*.

Let 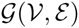 be a fully connected, undirected and non-reflexive graph where 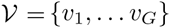 is a set of nodes (corresponding to genes), and 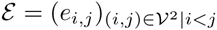 is a set of edges (corresponding to plausible gene interactions). Each edge *e_i,j_* is weighted by the value from matrix **W**. The initial number of gene-to-gene edges of 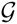, denoted by *ϵ*, is equal to *G*(*G* − 1)/2. Inferring a GRN from 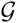 aims to construct a final graph selecting a subset of edges 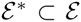 which reflects true gene regulatory processes. We formulate the search for this graph by computing an edge indicator vector **x** ∈ {0,1}*^ϵ^* whose components *x_i,j_* are such that

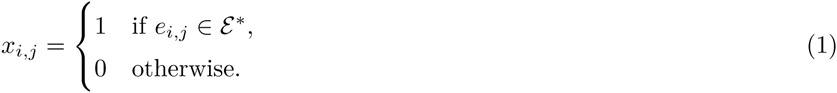

We assume in this work that a list of putative transcription factors is available. A gene supposed to code for a transcription factor is metonymically denoted by TF. A gene not identified with this property is designated by 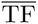. The 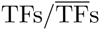 notation defines two complementary subsets of the ensemble of genes 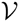. Subsequently, 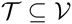 denotes the set of putative TFs. We consider that regulation is implicitly oriented from TF toward 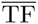 genes, and do not infer edge directions between TF-TF links. Assuming that significant edges have stronger weights *ω_i,j_*, we wish to maximize the sum of weights, while expressing our structural a priori in the inference model. To that goal, the edge labeling problem is formulated as the minimization of the composite functional:

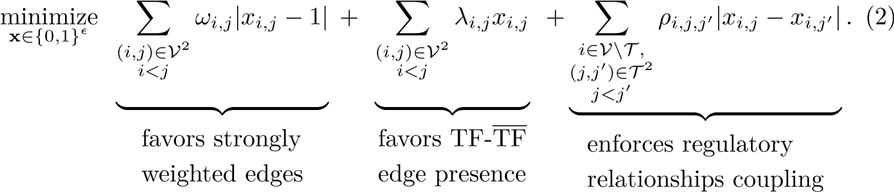

Let us comment the first term in the above functional. In order to select edges of strong weights *ω_i,j_*, the first term reflects a biological data fidelity term. It represents a gene-to-gene edge deletion cost. Thus, if *ω_i,j_* is large (respectively, small), its edge deletion cost is high (respectively, low), disfavoring (respectively, favoring) its deletion. We now explore the two last penalty terms of (2) corresponding to our biologically-related structural a priori regularization.

The second term counterbalances the first one. Independently from the fact that actual TF genes are less numerous than 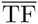 genes, regulatory relationships between couples of TFs are expected to be less frequent than between one TF and one 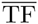. This expectation may promote biological graphs with a modular structure [19, 20]. An illustration is presented in Figure 1. As we are looking for gene regulatory knowledge, we infer edges linked to at least one TF. In addition, we want to favor the preservation of 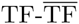 edges over TF-TF links. This edge selection capability is driven by positive weights *λ_i,j_*. Their values depend on the three types of pairs of nodes *i* and *j*. We define these case-dependent weights as follows:

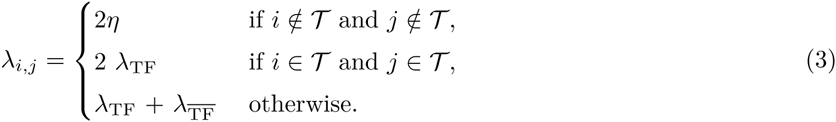

**Figure 1.**
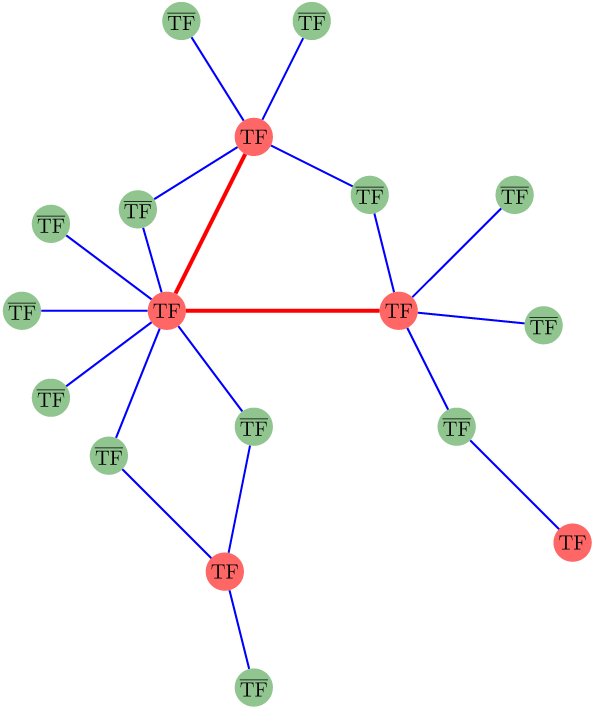
Illustration of our first a priori on a synthetic GRN. The TF-TF edges (red edges) are less represented than the 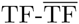 edges (blue edges). The red nodes code for the TF genes while the green nodes code for 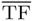 genes. The ratio of the number of TF on the total number of genes is 5/18 in this example. The ratio of the number of TF-TF edges on the total number of edges is 2/20, which is about 2.5 times smaller.

Hence, 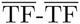 edges have weights assigned to 2*η*. The parameter *λ*_TF_ (respectively, 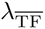) acts in the neighborhood of TF genes (respectively, 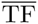 genes). They may be interpreted as two threshold parameters. This double threshold promotes grouping between strong and weaker edges among functionally-related genes. A similar approach is used in image segmentation [21] to enhance object detection with reduced sensitivity to irrelevant features [22]. To promote 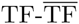 interactions, the λ_TF_ parameter should be greater than 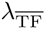. To ensure that any TF involved interaction is selected first, we should verify that 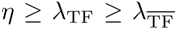. Additionally, removing all 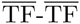 edges amounts to setting their corresponding *x_i_,_j_* to zero. Consequently, *η* should exceed the maximum value of the weights *ω*. Since we address different data types and input weight distributions, we can easily renormalize them all to *ω_i,j_* ∈ [0,1], and choose *η* = 1. When 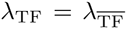, no distinction is made on the type of edges. This is equivalent to using a unique threshold value, as in classical gene network thresholding. This can be interpreted as if, without further a priori, all genes were indistinguishable from putative TFs. However, different λ_TF_ and 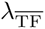 may be beneficial. We indeed show in the Supplementary Materials that for any fixed value of λ_TF_, smaller values for 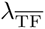 improve graph inference results. A simple linear dependence 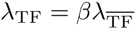, with *β* ≥ 1 suffices to define a generalized inference formulation encompassing the classical formulation. We fixed here *β* as a parameter based on the gene/TF cardinal ratio: 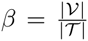. This choice is consistent when no a priori is formulated on the TFs (i.e. all genes are considered as putative TFs). Hence, *β* = 1 and 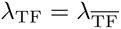. As mentioned above, without knowledge on TFs, we recover classical gene network thresholding. The *λ_i,j_* parameter now only depends on a single free parameter 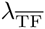, similarly to the large majority of inference methods requiring a final thresholding step on their weights.

Finally, the third term of the proposed functional aims to enforce a regulator coupling property (see Figure 2). If two transcription factors are co-expressed, and co-regulate at least one gene, we consider plausible that any gene regulated (respectively non regulated) by one of these TFs is regulated (respectively, non regulated) by the other TF. We quantitatively translate the co-expression of two TFs *j* and *j*′ by *ω_j,j_*_′_ > *γ*, where *γ* ∈ ℝ^+^ is a threshold reflecting the strength of the co-expression between *j* and *j*′. Similarly, the regulation of a 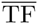 *k* by a TF *j* (respectively, *j*′) is numerically expressed by *ω_j,k_* > *γ* (respectively, *ω_j_*_′,_*_k_* > *γ*). We define *γ* from robust statistics [23] as the (*G* − 1)^th^ quantile of the weights. We thus choose the coupling parameter as:

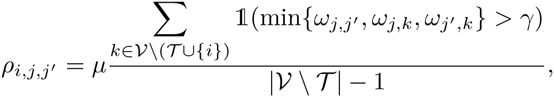

where 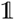 is the characteristic function (equals to 1 when the condition in argument is satisfied and 0 otherwise) and *μ* ≥ 0 is a regularization parameter controlling the impact of the third term on the global cost. The proposed numerator counts the number of 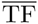 genes co-regulated by *j* and *j*′. As we exclude the gene i, the maximal number of TF genes co-regulated by *j* and *j*′ equals 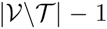. Hence, using the latter quantity as the denominator casts the *ρ_i,j,j_*_′_/*μ* parameter as a coregulation probability relative to couples of TFs (*j*, *j*′). The greater the co-regulation probability, the stronger the influence of the third term. This penalty requires that at least two 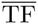 target genes exist (hence, the denominator does not vanish). Otherwise, when 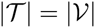, we set *μ* = 0.

**Figure 2.**
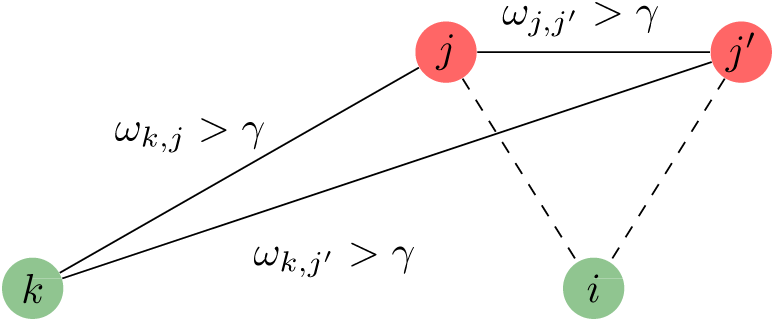
Regulator coupling property. If the transcription factors *j* and *j*′ are co-expressed (*ω_j,j_*_′_ is high, represented by a solid edge), and there exists at least one gene *k* that is not a TF, and is co-regulated by both *j* and *j*′, then the presence in the inferred graph of edge *e_i,j_* is coupled with the presence of *e_i,j_*_′_.

We now turn our attention to the strategy for computing an optimal labeling vector **x**^*^ solution to Problem (2).

### Optimization strategy

By using elements from graph theory, we now explain how a maximum flow algorithm can solve Problem (2). It relies on the maximum flow/minimum cut duality [24]: the computation of an optimal edge labeling minimizing (2) can be performed by maximizing a flow in a network *G_f_*.

A flow (or transportation) network 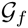 is a directed, weighted graph including two specific nodes, called source (a node with 0-in degree) and sink (a node with 0-out degree), respectively denoted by *s* and *t*. We recall that the degree of a node is defined as the number of edges incident to that node.

We now introduce the concept of flow in the transportation network 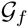. A flow function *f* assigns a real value to each edge under two main constraints: the capacity limit and the flow conservation. The capacity limit constraint entails that the flow in each edge has to be less than or equal to the capacity (i.e. the weight) of this edge. If the flow equals the capacity, the edge is said saturated. The flow conservation constraint signifies that, at each node, the entering flow equals the exiting flow. Subject to these two constraints, the aim is to find the maximal flow from *s* to *t* in the flow network 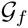. According to the graph construction rules provided by [25], the flow network for solving Problem (2) is composed of:

- A set of *ϵ* nodes *a_i,j_* with (*i*, *j*) ∈ (1, …, *G*}^2^, *i* < *j*, linked to the source *s* with edges of weight *ω_i,j_*. Each node is associated with a label *x_i,j_*.
- A set of *G* nodes *v_k_* of labels *y_k_* with *k* ∈ (1, …, *G*}. The nodes *v_k_* is linked to the previously defined node *a_i,j_* if *k* = *i* or *k* = *j*. If such an edge exists, a weight *λ_k_* is thus assigned. In reference to (3), the weight *λ_k_* equals *η*, *λ*_TF_ or 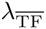, according to the nature of the node *a_i,j_* (corresponding to the edge *e_i,j_* in the initial network 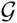).
- A set of *q* edges, linking nodes *a_i,j_* to *a_i,j′_* for which the regulator coupling property is satisfied, with weights equal to *ρ_i,j,j_*_′_.
- An additional set of *G* edges, linking nodes *v_i_, i* ∈ {1, …, *G*} to the sink node *t*.

Figure 3 illustrates this graph construction on a small-size example. Computing a maximum flow from the source to the sink in this flow network saturates some edges, thus splitting the nodes *a_i,j_* into two different groups: nodes that are reachable through a non saturated path from the source, and those that are not. Assuming that the source node *s* is labeled with 1, and the sink node *t* is labeled with 0, binary values are thus attributed to the edge labels *x_i,j_* (secondarily, binary values are also assigned to the *y* labels of nodes *v* in the flow network), and this final labeling returns the set of selected edges 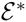 which minimizes (2). We use the C++ code implementing a max-flow algorithm from [26].

### Problem dimension reduction

As explained in the previous section, the optimal solution to the minimization problem (2) may be obtained *via* a maximum flow computation in a network generated from the whole original graph. In practice, many parameters *ρ_i,j,j_*_′_ have zero values. So rather than building 0-valued edges in the flow network, reducing the dimension of this network is judicious. Indeed, if *ρ_i,j,j_*_′_ = 0 for all 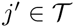, the optimal label of *x_i,j_* is given by the explicit solution

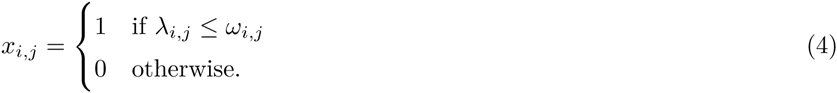

This formula also provides a better insight into the role played by thresholding parameters *λ_i,j_*. We now have a fast optimization strategy to generate a solution to the proposed variational formulation. One of the advantages of employing the BRANE Cut algorithm is the optimality guaranty of the resulting inferred network with respect to the proposed criterion. We next describe quantitative gains that can be achieved using BRANE Cut.

**Figure 3.**
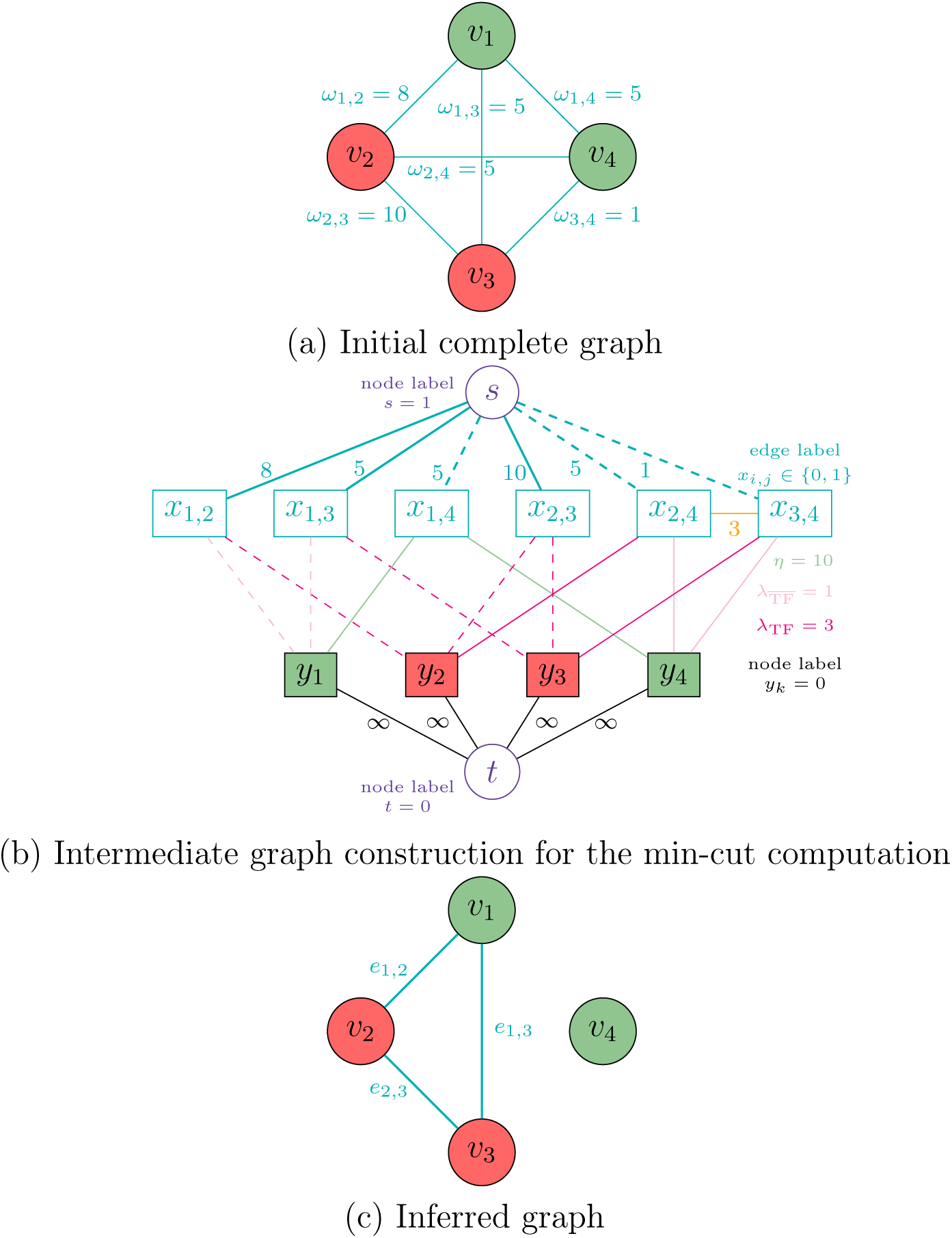
Schematic view of the proposed BRANE Cut method. The initial graph (a) is transformed into an intermediate graph (b) in which a max-flow computation is performed to return an optimal edge labeling **x**\* leading to the inferred graph (c). We choose to present the method in its full generality with unscaled weights (i.e. *w_i,j_* ∈ [0, +∞], and *λ* parameters also belong to [0, +∞[). Nodes *v*_2_ and *v*_3_ are TFs, 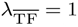 and *λ*_TF_ = 3. Taking *γ* = 4 implies that *v*_1_, *v*_2_, and *v*_3_ satisfy the regulator coupling property. Vertices *v*_1_ and *v*_4_ are thus affected, leading to the presence of additional edges weighted by *ρ*_1,2,3_ = 0 and ρ_4,2,3_ = 3, when *μ* is set to 3. Computing a max-flow in the graph (b) leads to some edge saturation, represented in dashed lines. The values from the source (value 1) and the sink (value 0) are propagated through non saturated paths, thus leading to *x*_2,4_ = *x*_3,4_ = 0.

## Results and Discussion

We compare the BRANE Cut approach to the top performing graph inference methods on synthetic and real data. The considered state-of-the-art methods are CLR, which outperforms ARACNE and Relevance Networks on the *E. coli* dataset, and GENIE3, winner of the DREAM4 multifactorial challenge [27] on synthetic data among a large number of competing methods, and also outperforming other approaches on the real *E. coli* dataset. For a fair evaluation, all networks are inferred using the same set of parameters for a given method: CLR results are computed with the ’plos’ method and the default values for the two quantization parameters. GENIE3 outcomes are obtained using the Random Forest method and 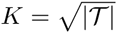. We also postprocessed both the CLR and GENIE3 weights with ND (network deconvolution [12]), and applied BRANE Cut on both the deconvolved ND-CLR and ND-GENIE3 networks.

### Validation datasets

#### The DREAM4 dataset

The Dialogue for Reverse Engineering Assessments and Methods fourth (DREAM4) [27] multifactorial challenge provides five simulated datasets with real network topologies from the prokaryote *E. coli* and the eukaryote *S. cerevisiae*, and simulated expression data. At the time of the challenge, the competing approaches did not have access to a list of putative transcription factors, which is now available online. As this list is a requirement of our method, we benchmark the best performing network inference methods using this additional information. The networks are composed of 100 genes, with a total of 100 expression levels per gene. The evaluation of the inferred networks was performed using the gold standard provided in the DREAM4 multifactorial challenge.

#### The Escherichia coli dataset

This dataset was first introduced in [8] and is composed of 4345 gene expression profiles, each profile containing 445 gene expression levels. This compendium contains *steady-state* and *time-course* expression profiles. RegulonDB [28] is the primary database on transcriptional regulation in *Escherichia coli* K-12 containing manually curated knowledge from original scientific publications. As in [8], we used the version 3.9 to evaluate inferred networks. This database offers a set of 1211 genes for which 3216 regulatory interactions are confirmed.

#### The DREAM5 dataset

The DREAM5 challenge (Dialogue for Reverse Engineering Assessments and Methods fifth) [2] provides four networks. The first one contains an in-silico dataset while the three others correspond to real datasets. For the four networks, the list of putative transcription factors is known. In this work, we used the three networks (1, 3 and 4) for which a ground truth is provided. The first network is composed of 1643 genes (195 TFs) and expression data in 805 conditions. Network 3 contains information about 4511 genes (334 TFs) in 805 conditions, while network 4 compiles 5950 genes (333 TFs) and 536 conditions. The evaluation of the inferred networks was performed using the gold standard provided in the DREAM5 challenge.

### Evaluation measures

Predictive measures, standard in binary classification or machine learning, benchmark different network inference methods. For a given network, Precision and Recall (sensitivity) are defined as

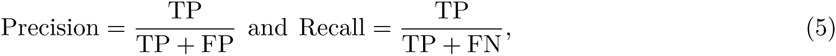

where TP is the number of true positive, FP is the number of false positive and FN is the number of false negative. The Precision value indicates the proportion of correctly inferred edges compared to the total number of inferred edges. The Recall value reveals the proportion of correctly inferred edges compared to the total number of expected edges given by the gold standard. In order to evaluate and to rank the different tested methods, Precision-Recall (PR) curves are commonly used [8]. As the best results correspond to both high precision and high recall values, the Area Under the Precision-Recall Curve (AUPR) is an appropriate quantitative criterion to measure the quality of an inference method (higher is better).

### Results on DREAM4

To validate our BRANE Cut approach, we used a variety of different initial weights, directly obtained from CLR, GENIE3, or after ND postprocessing [12] (ND-CLR and ND-GENIE3). Similarly to BRANE Cut, ND takes weights given by other inference approaches as inputs. When necessary, input weights are symmetrized by retaining the maximal value between *ω_i,j_* and *ω_j,i_*. The comparison of each generated graph to the ground truth for each network allows the construction of five Precision-Recall curves. They are obtained from all the different possible threshold 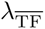 values and are provided in the Supplementary Materials. All networks are generated setting *μ* = 3 and *γ* takes the (*G* − 1)^th^ quantile value of the normalized weights *ω*. Quantitative results are reported in Table 1. We provide a heuristic to determine *μ* and perform its sensitivity analysis in the Supplementary Materials.

Computed AUPR in Table 1 (a) highlight in bold that, globally, first and second best performances are always produced with BRANE Cut. Furthermore, each method tested (CLR, GENIE3, ND-CLR or ND-GENIE3) used as initialization exhibits an improved AUPR with BRANE Cut post-processing. Indeed, the average improvement reaches 10.6% based on the CLR weights, 8.4% for the GENIE3 weights, 5.9% with ND-CLR weights and 7.2% compared to the ND-GENIE3 weights, see Table 1 (b).

We finally compare ND and BRANE Cut as post-processing methods on original weights. As shown in Table 1 (c), BRANE Cut outperforms network deconvolution except for a practically unnoticeable degradation on the fifth network for GENIE3 weights. This relative improvement is essentially due to the fact that network deconvolution degrades results on the first two networks.

In the associated Precision-Recall curves, reported in the Supplementary Materials, we notice that the improvements of our results are mostly obtained in the first part of the curves, corresponding to a Precision greater than 50% in the inference. Thus, such inferred graphs are expected to be more reliable for a biological interpretation. From this observation, looking at the AUPR for different Precision ranges, from the whole scale to precisions above 50%, provides a finer assessment of the predictive power of inference methods. Thus, Figure 4 highlights relative AUPR improvements for given Precision ranges. It illustrates that BRANE Cut improvement ratios over GENIE3 AUPR are clearly visible at higher Precision ranges, typically over 65%.

**Table 1.**
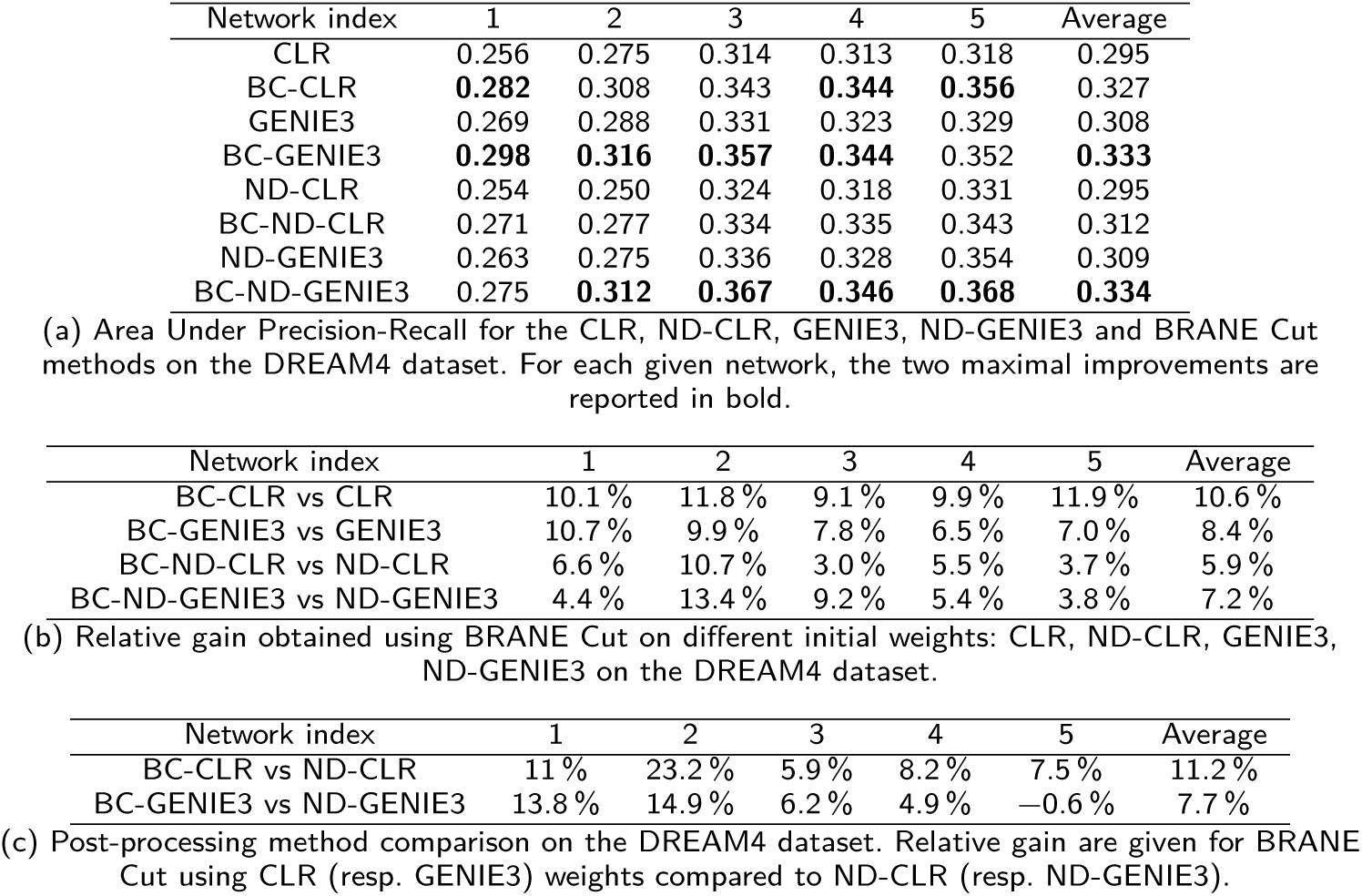
BC-X corresponds to the BRANE Cut method initialized with the weights of the method X.

**Figure 4.**
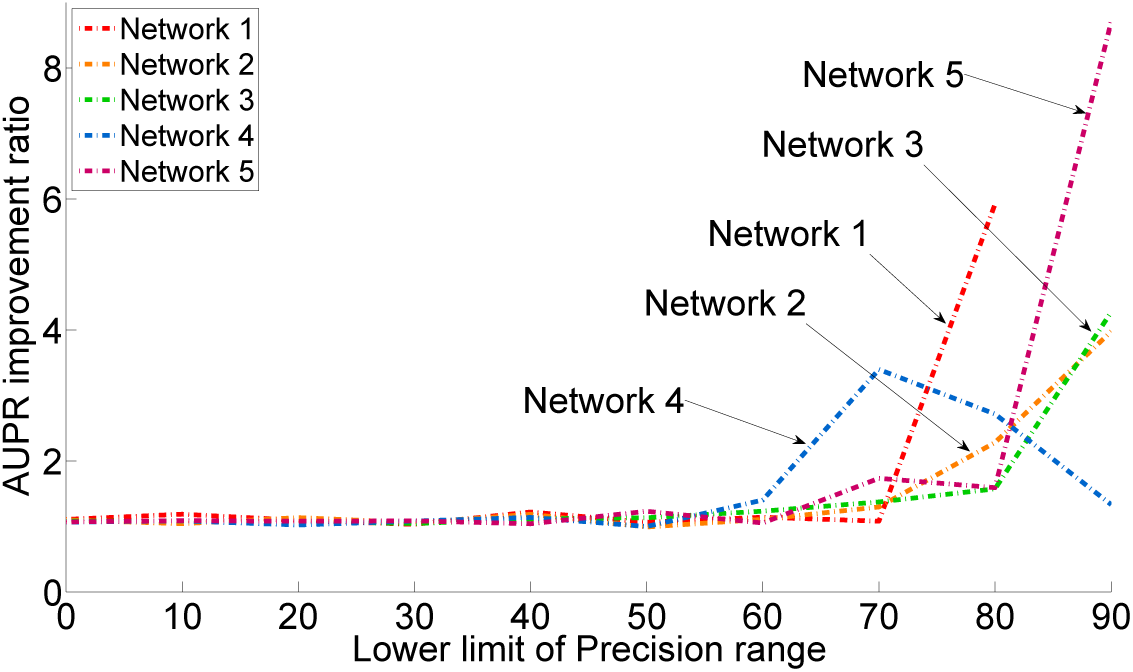
AUPR improvements for different parts of the PR curves on the five networks of DREAM4. In order to show the differential improvement over the Precision, relative AUPR are computed for different PR curves, truncated at different range of Precision: [0,100], [10,100], …, [90,100]. Here, the improvement is defined as the AUPR ratio of BC-GENIE3 and GENIE3.

Based on the AUPR criterion, we conclude that BRANE Cut outperforms state-of-the-art methods. Specifically, single-threshold results are sensibly refined by our approach, regardless of initial weights.

### Results on the *E. coli* dataset

We now present the results of the BRANE Cut method on the real *E. coli* dataset. Our approach uses the normalized weights *ω_i,j_* defined by CLR (with *μ* = 1000) and GENIE3 (with *μ* = 10). A discussion about the choice of the *μ* parameter is given in the Supplementary Materials. The parameter *γ* is set as in the previous section. The different Precision-Recall curves are reported in Figure 5, to compare BRANE Cut to GENIE3 and CLR.

**Figure 5.**
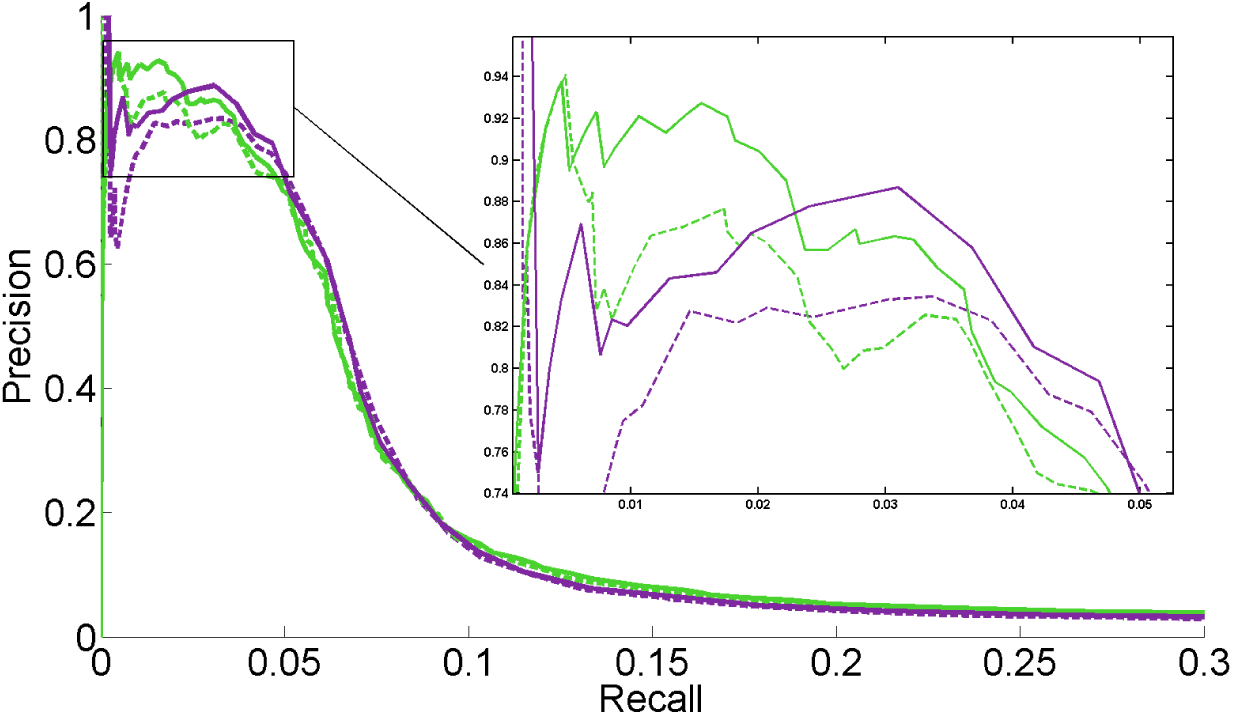
Precision-Recall curve comparison on the *E. coli* dataset. CLR (dashed purple line) and GENIE3 (dashed green line) are compared to our BRANE Cut method initialized with the weights CLR (solid purple line) or GENIE3 (solid green line).

Best performance is expected toward the upper right side of Precision-Recall curves, with both high recall and precision. However, GRN Precision-Recall curves traditionally exhibit low Precision values over the whole curve on real datasets due to the difficulty in inferring accurate regulation relationships among large amounts of genes. For instance with the *E. coli* dataset, we observe that a recall below 0.05 corresponds to small inferred graphs, with less than 300 edges and a high precision (more than 75%). Due to their higher predictive power and their readability, such small networks are often preferred by biologists. Hence, we focus on the upper-left part of the Precision-Recall curves in Figure 5, emphasized in a close-up, corresponding only to high precision and small graphs. Here, BRANE Cut initialized with GENIE3 weights proves to be the best performer on smaller graphs (less than 100 edges corresponding to a recall below 0.02). However, graphs of larger size (up to a recall of 0.08) are more accurately reconstructed with CLR and BRANE Cut initialized with CLR weights. Again, the BRANE Cut approach improves the prediction results of both CLR and GENIE3.

Overall, as reported in Table 2, BRANE Cut produces better results in terms of AUPR. Specifically, relative gains presented in Table 2 show a significant enhancement of CLR results and a more moderate enhancement of GENIE3 results. Taking into account that CLR weights are obtained more than seven times faster than GENIE3 weights, BRANE Cut initialized with CLR weights finally recovers results comparable to those obtained by GENIE3, but much faster. Initializing BRANE Cut with the GENIE3 weights, the results are still improved with negligible additional times compared to weight computation.

**Table 2.** Area Under Precision-Recall, computation times and relative gains on the *E. coli* dataset using BRANE Cut with CLR or GENIE3 weights.

|  | CLR | BC-CLR | GENIE3 | BC-GENIE3 |
| --- | --- | --- | --- | --- |
| AUPR ($\times 10^{-2}$) | 7.86 | 8.79 | 8.90 | 9.17 |
| Total comput. time (min) | 41.0 | 41.05 | 303 | 303.05 |
| Gain | 11.8% AUPR gain over CLR<br>7.4 × faster than GENIE3 | | 3.0% AUPR gain over GENIE3<br>negligible additional computation cost | |

Table 3 shows network inference improvements using BRANE Cut in terms of the number of verified inferred edges for comparable Precision values.

**Table 3.** Comparison of graph inference in terms of number of True Positive edges and Recall at constant Precision using GENIE3 or BRANE Cut-GENIE3 on the *E. coli* dataset.

| Precision (%) | Recall (%) |  |
| --- | --- | --- |
|  | GENIE3 | BRANE Cut |
| 90 | 0.55 | **2.00** |
| 85 | 2.31 | **3.40** |
| 80 | 3.77 | **3.86** |
| 75 | 4.19 | **4.55** |
| Precision (%) | TP edges |  |
|  | GENIE3 | BRANE Cut |
| 90 | 18 | **66** |
| 85 | 75 | **112** |
| 80 | 124 | **127** |
| 75 | 138 | **150** |

#### *Inferred network example on* E. coli

An example of regulatory network on the *E. coli* dataset obtained with BRANE Cut, initialized with GENIE3 weights, is displayed in Figure 6. The inferred network obtains a Precision score of 85%, with a better predictive power than the network produced by the GENIE3 method alone. The binary network for GENIE3 is obtained by selecting edges having a weight higher than 0.707. This threshold renders a network with 85% of Precision. In comparison to the reference, we discover 20 additional plausible regulatory interactions. Among these 20 predictions, ten were also predicted by the GENIE3 method, leading to ten predictions specific to BRANE Cut. By analyzing the predictions using STRING [29] and EcoCyc [30] databases, we observe that the predicted groups of genes were already identified as co-expressed and are known to belong to the same functional mechanism.

**Figure 6.**
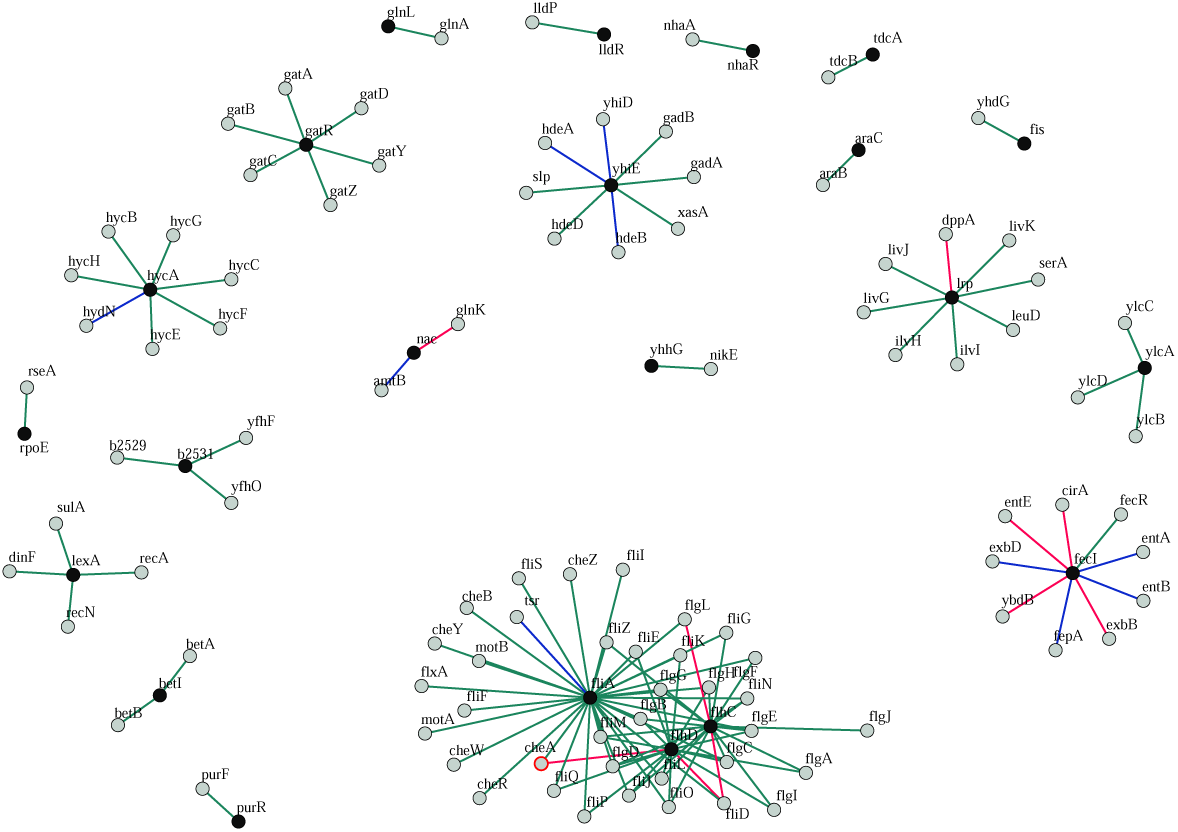
Example of network built using BRANE Cut on the *E. coli* dataset. Legend: black nodes: transcription factors, gray nodes: other genes. green edges: inferred regulations also reported in the gold standard, blue edges: new inferred regulations that are also inferred by GENIE3, and pink edges: new inferred regulations.

#### Influence of the proposed structural a priori

We start to analyze the influence of our first a priori on the *E. coli* dataset using CLR weights. Hence, using the first two terms with 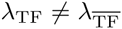 leads to an AUPR of 0.0870, which constitutes a relative improvement of 10.7% over CLR, without co-regulation a priori. More generally, as *λ*_TF_ and 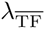 are interpreted as a pair of thresholds, the higher these parameters, the greater the stringency in the inferred graph. These results show that allowing a different threshold value in the neighborhood of transcription factors than for other genes does play a positive role by itself. The regulator coupling term controlled by *μ* brings further improvements. Indeed, the addition of the third term results in an AUPR equal to 0.0879, corresponding to a relative improvement of 11.8% over CLR. The corresponding Precision-Recall curves are displayed in the Supplementary Materials. They show that even if the gain brought by the co-regulation a priori is shallower than the improvement allowed by the first a priori, it remains valuable despite its localization in the high Precisions area.

#### Algorithmic and computational complexity

As previously mentioned in the Optimization strategy section, we used the C++ code implemented by [26]. Using this algorithm, the computational complexity of BRANE Cut is *O*(*mn*^2^|*C*|), where *m* (respectively *n*) is the number of edges (respectively the number of nodes) in the flow network 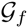, and |*C*| the cost of the minimal cut. Specifically, in our case (without the dimension reduction trick presented in the Problem dimension reduction section) the number of nodes in the flow network is equal to the sum of the number of edges *ϵ* in the initial network, the number of genes *G* plus two additional nodes (source and sink). The number of edges *n* is equal to 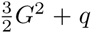, where *q* is the number of edges coding for the co-regulation a priori. Note that, as mentioned in [26], this complexity is not the best achievable by a max flow algorithm. Meanwhile, their experiments showed better performance for several typical computer vision problems. Not being in a computer vision setting, we could benefit from faster max flow algorithms. However, since the time spent on max flow computation to infer the large graph of *Escherichia coli* is small (only several seconds), the benefit would not be noticeable.

Given pre-computed weights, our algorithm requires 30 additional seconds to infer the *E. coli* network, without using the simplification described in the Problem dimension reduction section. By computing the explicit solution to our problem on a subset of edges, we improve BRANE Cut computation times by a factor of 10. Given CLR weights computed in 41 minutes on a Intel Core i7, 2.70 GHz laptop, our algorithm thus only requires three additional seconds. We note that the weight computation duration of GENIE3 are sensibly longer (5 hours), using the list of transcription factors. If one wished to build a *E. coli* network that would also contain 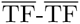 interactions using GENIE3, it would take 20 minutes per gene, for a total of two months with a basic rule of three.

### Results on DREAM5

We have evaluated BRANE Cut on three DREAM5 networks (1, 3 and 4) for which a validation exists. BRANE Cut parameters are initialized with the proposed heuristics and results are obtained using the validation procedure previously detailed. BRANE Cut outperforms CLR and GENIE3 by 7% and 5% respectively on Network 1. The improvement is 2.8% and 2.1% for Network 3. For the fourth network, the maximum Precision only reaches about 35%. The AUPR computed with every method is exceptionally low. As such, the relative AUPR differences are insignificant, within the numerical precision. The detailed AUPR are given in the Supplementary Materials. Regarding the results in these additional datasets, the proposed heuristics lead to improvements over state-of-the-art.

## Conclusions

By using structural a priori that are often available but rarely used, we managed to infer networks that recover more true interactions than previous approaches, on both synthetic and real datasets. We have expressed the graph inference as an optimization problem, and used the generic Graph cuts approach, very popular in computer vision, to compute the optimal edge labeling of our inferred graph. Comparisons are performed with simple regularization parameters based on gene set cardinality. We obtain better results than both CLR and GENIE3 in terms of Area Under the Precision-Recall curves, even with ND deconvolved networks. BRANE Cut yields state-of-the-art results, with a negligible computation time. While the GENIE3 method needs about five hours to obtain a 4345-gene network, limited to interactions involving transcription factors, we obtain a network with similar accuracy with our method in a few seconds, only using CLR weights computed in about forty minutes. This graph inference acceleration is thus useful to explore large datasets. Some predictions specifically identified by our method indeed appear as relevant interactions.

As mentioned in [2], community-based methods represent a promising future for gene network inference, by aggregating the predictions of existing GRNs approaches. As our method takes any weights as an input, it has the potential to improve other GRN approaches providing pairwise weights. Results provided in the Supplementary Materials illustrate these remarks.

Based on these assessments, a perspective consists of the integration of various weights, provided by competing GRN methods, to further improve and strengthen present results. This integration may involve multi-valued graphs or network fusion [31].

### Authors’ contributions

AP participated to the mathematical modeling, implemented the software and drafted the manuscript. CC co-designed the mathematical modeling, suggested the solver and took part to the writing of the paper. LD participated to biological data processing and performance evaluation, manuscript writing, and contributed to the bibliographic study, FB participated to the biological a priori definition and to the discussion of the results in the manuscript. JCP co-designed the mathematical modeling, participated to speedup improvements and to the manuscript writing. All authors read and approved the final manuscript.

## Acknowledgements

We thank V. A. Huynh-Thu for her help in our comparisons with GENIE3, and Bruno Léty for careful proofreading. We are indebted to the reviewers for their insightful remarks which contributed to significantly improve the quality of our paper.

## Additional Files

Additional file — Supplementary Materials

The supplementary file provides detailed justification for the choice of the model parameters, studies their relative influence and provides a sensitivity analysis.

